# Effects and mechanisms of monoclonal and polyclonal human antibodies in protection of humanized mice from HIV-1 challenge

**DOI:** 10.1101/2025.11.12.688003

**Authors:** Urjeet S. Khanwalkar, Jennifer Fields, Joshua A. Wiener, Marissa L. Lindman, Alejandro B. Balazs, Michael S. Seaman, Lindsey R. Baden, Stephen R. Walsh, Steven N. Fiering, Yina H. Huang, Margaret E. Ackerman

## Abstract

Recent clinical trials of both active and passive immunization demonstrate the barriers to the successful development of efficacious preventative HIV-1 vaccines and prophylactic antibody treatments. More facile means to explore these interventions in preclinical models could fill key gaps in knowledge and contribute to identifying and optimizing the most promising interventions. Here we report simple adaptations to standard virus challenge strategies in the humanized mouse model that support evaluation of antibody-mediated protection from infection with fewer high stakes design choices and demonstrate evaluation of protection afforded by polyclonal human serum IgG antibodies (pAbs) and monoclonal antibodies (mAbs) with variable pharmacokinetic (PK) and functional profiles. Using these adaptations, we observed that both neutralization and Fc-mediated effector functions contribute to the *in vivo* antiviral activity of broadly-neutralizing antibody VRC01, that confounding of results due to differences in mAb PK can be overcome, and most promisingly, that polyclonal human serum IgG that exhibits potent neutralizing and Fc-effector function can protect from infection. Collectively, this work demonstrates insights into antibody-mediated protection and methods that hold promise in supporting testing the protection from HIV-1 afforded by human pAb responses induced by vaccination.

## Introduction

After favorable results of the RV144 HIV-1 vaccine trial were reported in 2009^1^ and offered hope to the field, challenges for the development of a successful HIV vaccine have become more apparent. Multiple major efficacy trials of active vaccination regimens^2–4^, as well as the first clinical efficacy trials of passive immunization with a broadly neutralizing antibody^5^ failed to meet their predefined primary efficacy criteria. However, follow-up studies identified both virological and immunological correlates of risk (CoR) in case-control study designs^5–10^ that suggested that some individuals were protected from some viruses by these interventions. More facile means to probe these associations in preclinical models and better define their mechanistic relevance would fill key gaps in knowledge about how to both identify and optimize promising interventions to prevent HIV-1 infection.

To this end, preclinical models that mimic human HIV infection are essential for evaluating novel therapeutics. Among these, the bone marrow-liver-thymus (BLT) humanized mouse model^11–13^ provides a substantial advance by offering a small animal alternative to nonhuman primate (NHP) models. Humanized mouse models offer practical advantages of increased power and accessibility to investigators outside of dedicated primate centers, and biological advantages of human target and effector cells and authentic virus rather than simian immunodeficiency virus or simianized HIV. The BLT model involves co-transplantation of human fetal liver and thymic tissues along with CD34+ hematopoietic stem cells into immunodeficient NSG (NOD.Cg-Prkdc^scid^ Il2rg^tm1Wjl^) mice. The BLT model supports robust human T cell development and is highly permissive to HIV infection, making it invaluable for studying viral pathogenesis and immune responses. Similarly, the CD34+ humanized mouse model^14,15^, which relies on the engraftment of CD34+ human stem cells alone, results in partial reconstitution of the human immune system, providing similar support for HIV infection studies. While these models have been enabling, notable limitations remain, including incomplete immune system development, variability in engraftment efficiency, and imperfect human cell functionality.

Improvements afforded by use of modified mouse backgrounds include improved NK cell reconstitution^16,17^ observed on the NSG-SGM3 background (S15), better neutrophil reconstitution achieved in MISTRG mice^18^, and more functional adaptive immune responses in the humoral compartment afforded by the new THX (17β-estradiol-conditioned Kit^W-41J^) model^19^. These models continue to further our understanding and advance promising preventative as well as therapeutic interventions. Collectively, they have contributed to studies of viral latency^20^, small molecule antiretroviral drug therapy (ART)^21^, complex kick and kill functional cure strategies^22^, and chimeric antigen receptor-based interventions^23^, among others. Beyond testing the efficacy of different interventions, they have provided critical insights into the mechanisms of viral control, resistance, and immune-mediated clearance, thus accelerating the translation of promising therapeutic strategies.

In particular, humanized mouse models have been effectively used to better understand the antiviral effects of broadly neutralizing antibodies (bNAbs) *in vivo*. They have been used to evaluate the prospects of persistent bNAb expression to prevent infection^24^, of individual and bNAb combinations to effectively suppress established infection^25^, and of bNAbs and viral inducers to decrease viral rebound^26^. Antiviral activities have been mechanistically probed in studies evaluating bNAb variants with ablated or improved antibody effector function in models of both prevention and therapy^27–30^—covering a range of antibodies and infection states that would have been prohibitively resource-intensive to explore exclusively in NHP models. Similarly, the potential mechanistic relevance of antibodies that lack potent or broad neutralizing activity have been tested^31^, and modulation of these activities by viral genes such as Vpu, or by co-treatment with CD4 mimetics that change the conformational state of envelope glycoprotein have been determined^32^. Each of these examples present experimental questions that were answered leveraging designs made more practically accessible in humanized mice.

Nonetheless, challenges remain. Here, we focus on those that impact studies seeking to evaluate protection from infection in the context of a controlled challenge, which, unlike therapeutic interventions, cannot be ethically modelled in the clinic. As in NHP experiments, the viral dose used in challenge studies must be finely calibrated to the antibody dose so as not to either overwhelm any protective effects or fail to differentiate between antibodies that offer differing degrees of protection. A miscalculation in this aspect of design, whatever its experimental origin, can easily prevent a study from generating data addressing the primary hypothesis^33^. Differences in challenge and antibody dose may also alter the mechanism of action—a phenomenon hypothesized to exist for HIV^34^, and observed for other viruses^35^. Similarly, the time interval between administration of antibody and challenge is typically optimized to result in peak drug levels at the site of challenge, but these levels vary among animals in both tissue as well as in serum and can also vary from antibody to antibody. Each of these factors introduces experimental variability to the system—reducing the ability for models to finely discriminate between more and less effective interventions.

Additionally, the more rapid clearance of antibodies in mice than in NHP poses challenges to the repeat, low-dose challenge models favored as reflecting human exposure conditions. Indeed, repeat challenge of humanized mice has been used primarily in the context of vectored antibody delivery^29^, in which antibody is continually expressed by host cells rather than administered as recombinant protein. Distinct clearance rates and peak drug levels following one-time antibody dosing, which appear to be commonly observed for antibodies with modified Fc domains^33,36–39^, can greatly confound experiments in the NHP model^39^. Given the strong dependence of protection on antibody dose^29^, pharmacokinetic (PK) differences are likewise expected to confound results in humanized mouse models. Antibody can be dosed repeatedly, but this adaptation comes with a higher material cost and may nonetheless leave systemic and mucosal sites with varying drug levels that may not meaningfully reflect differences relevant in the clinic. Fortunately, the increased demands for drug are more practically accessible at humanized mouse than NHP scale, as seen from the prevalence of studies incorporating this feature^25–27,31,32^.

One area in which it is less feasible to increase dose is in passive transfer of polyclonal, serum-derived pools of antibody. Several studies have conducted passive transfer of polyclonal serum IgG^40–46^. However, these undertakings represent heroic efforts to fractionate liters of serum that, with one possible exception^40^, have more often than not yielded negative or only marginally positive results that are difficult to interpret. In other settings in which antibody quantities were limited, information about direct effects on virus particles or at mucosal sites was desired, or PK profiles were unknown or potentially prohibitive of systemic administration, investigators have employed mucosal administration of antibody shortly before challenge^47–51^ or mixed antibody with the virus inoculum preceding challenge^35^. Less materially intensive methods to evaluate the antiviral effects of antibodies *in vivo* such as these could be transformative to the study of the activity of polyclonal antibodies (pAbs) against HIV. For example, the breadth and degree of protection afforded by vaccine-induced antibodies could be evaluated *in vivo*—ideally in experiments that do not require more than the typical sample volumes banked in even early phase I trials.

Here, with this goal in mind, we explored use of adaptations of challenge strategies commonly used in NHP studies to reduce biological and technical variability and evaluate the mechanisms of antibody-mediated protection from HIV in the humanized mouse model. These modifications include escalating viral challenge dose so as to reduce the risk of wholly uninformative experiments; repeat virus challenges so as to improve power in the ability to dissect differences between treatment groups of limited size; and pre-mixing virus and antibody preparations as a means to reduce variance associated with drug PK and to make passive challenge experiments with limited antibody, such as pathogen-specific human pAb present in vaccine recipient serum, more feasible.

## Results

### VRC01 IgG1 demonstrates dose-dependent protection from mucosal HIV challenge

We first established the ability of neutralizing antibody to protect from infection when pre-mixed with virus as an intravaginally-administered inoculum, and to determine whether this approach was sufficient to show dose-dependent differences in protection. Accordingly, two independent batches of NSG-BLT humanized mice (n = 34) were challenged every two weeks with an increasing (3, 9, and then 27 ng of p24) amount of transmitted founder virus (REJO.c) that was incubated with low, moderate, or high (25, 100, or 400 μg) dose VRC01 or a high (400 μg) dose of CH42, a herpes simplex virus-specific isotype control antibody, for one hour prior to use as inoculum in an intravaginal challenge (**Figure 1A**). For this and all *in vivo* experiments, flow cytometry of peripheral blood cells was used to confirm suitable degrees of humanization based on frequencies of human CD45^+^ and CD4^+^ cells (**Supplemental Figure 1**), and groups were balanced for these cell distributions. Infection was monitored by RT-qPCR analysis of blood drawn weekly and time of infection was defined as the first timepoint at which plasma viremia above the limit of detection was observed.

**Figure 1.**
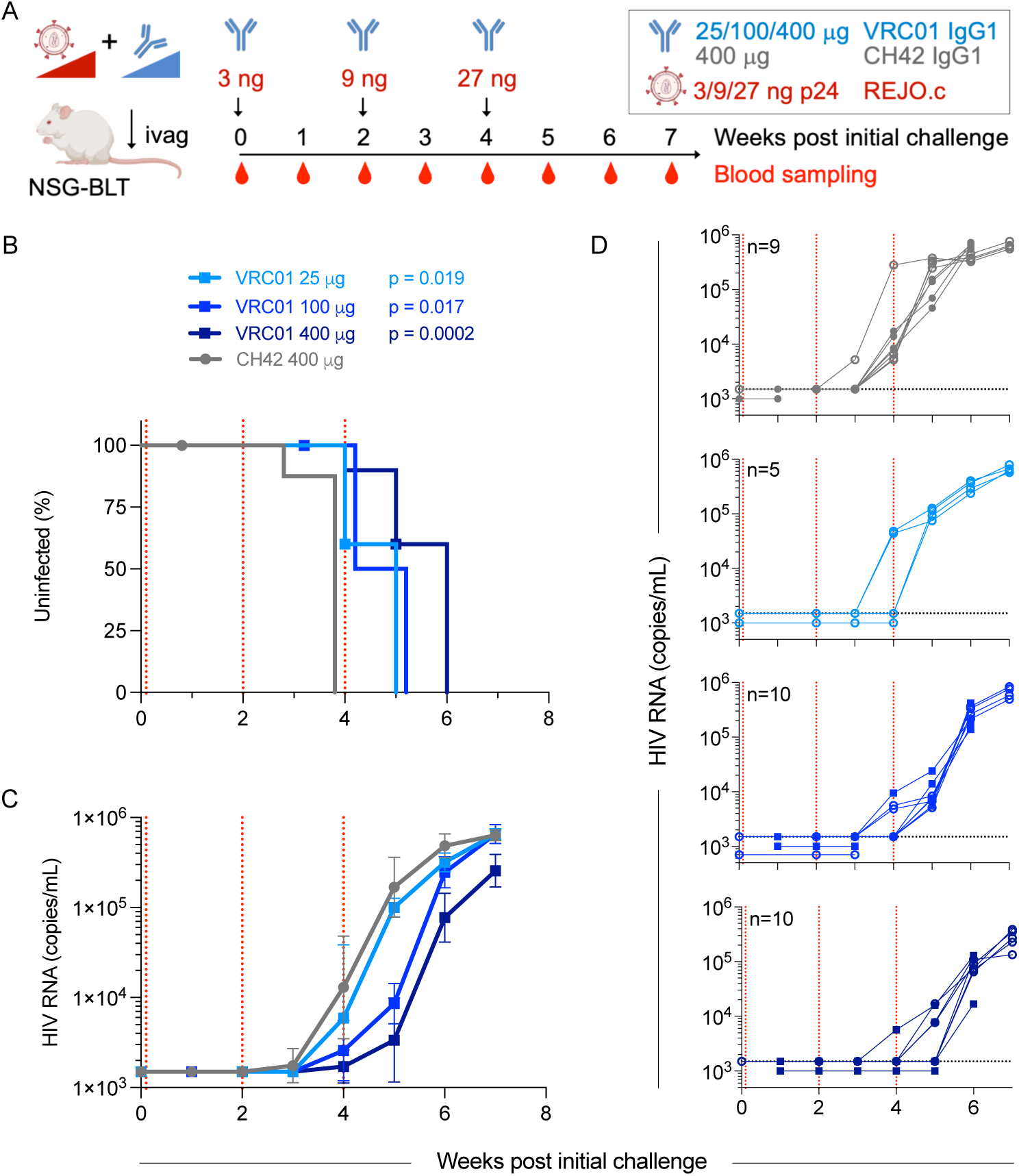
Efficient demonstration of dose-dependent protection from vaginal challenge in humanized mice by VRC01. **A.** Study design. Two batches (filled and hollow symbols) of NSG-BLT mice were serially challenged intravaginally (ivag) with a cocktail of REJO.c virus and mAb as illustrated. **B**. Kaplan-Meier curve of infection over time. Symbols indicated censored animals. **C-D**. Viral RNA levels over time. Geometric mean and 95% CI of viral RNA over time in each treatment group (**C**) and individual animals (**D**). Dotted horizontal line indicates limit of detection. Dotted vertical lines indicate virus challenge. Animals per group indicated in inset. Statistical significance of VRC01 treatment compared to isotype control defined by log-rank test.

Whereas no animals were infected following the initial, low-dose virus challenge, plasma viremia was observed in all animals that received the isotype control inoculum following the second, moderate-dose virus challenge by the day of the third, high-dose virus challenge (**Figure 1B**). In contrast, approximately half of each group of animals treated with VRC01 were aviremic at the time of the third challenge (**Figure 1B**). The low and moderate (25 and 100 μg) VRC01 dose groups showed similar rates of infection over time (**Figure 1B**), but infected animals in the moderate antibody dose group exhibited lower levels of viremia at earlier timepoints post-infection than those in the low antibody dose group (**Figure 1C,D**). All mice in these groups were infected by one week after the third (high dose) challenge, whereas only 30% of animals in the high (400 μg) VRC01 dose group were infected at this timepoint (**Figure 1B-D**). As compared to isotype control, VRC01 at any dose provided significant protection from infection (p ≤ 0.017 by log-rank test). Greater protection was conferred by high dose VRC01 than either moderate or low antibody doses (p = 0.0053 and p = 0.038, respectively). Animals in both batches of mice displayed similar susceptibility to infection and degrees of protection offered by the tested doses of broadly neutralizing antibody (**Figure 1D**). Collectively, these data demonstrate that in the pre-mixing model, both viral dose-dependent rates of infection and antibody dose-dependent protection can be reproducibly observed.

### 10-1074 IgG1 Fc variants with distinct functional and pharmacokinetic profiles

Variable PK observed both between animals as well as between antibodies contribute noise to preclinical models in which antibody is dosed prior to virus challenge, as different antibody levels are present in serum and tissue at the time of virus exposure. In studies evaluating repeated challenges following single antibody doses, varying antibody clearance rates over time can substantially impact the degree of protection observed^52^. When insights into mechanism of action and degree of protection rather than dose-dependence are desired, variable PK poses a significant challenge to experimental design and interpretation^33,39^. We next directly evaluated the ability of the pre-mixing model to exhibit reduced variance in association with antibody PK. To accomplish this goal, we used a panel of Fc-engineered variants of the broadly neutralizing antibody 10-1074. In addition to unmodified IgG1, an Fc-effector function knockout variant (LALA, L234A/L235A^53^), and a complement deposition-enhanced variant (EG, E430G^54^) with rapid clearance in NHP^39^ were generated. These antibodies demonstrated equivalent binding to gp120 TRO envelope protein (**Figure 2A**), and equivalent neutralization of REJO.c virus in the TZM-bl assay (**Figure 2B**). In contrast, the CH42 IgG1 isotype control showed no binding or neutralization activity. Fc variants were evaluated in a panel of *in vitro* assays of effector function (**Figure 2C-F**). Whereas activity of 10-1074 LALA was reduced in all assays, 10-1074 EG showed the expected increase in C3b deposition (**Figure 2C**) and complement-mediated viral lysis (**Figure 2D**), but no enhancement of phagocytosis (**Figure 2E**), or FcγRIIIa signalling (**Figure 2F**), which was assessed as a surrogate for NK cell-mediated antibody-dependent cellular cytotoxicity (ADCC) activity.

**Figure 2.**
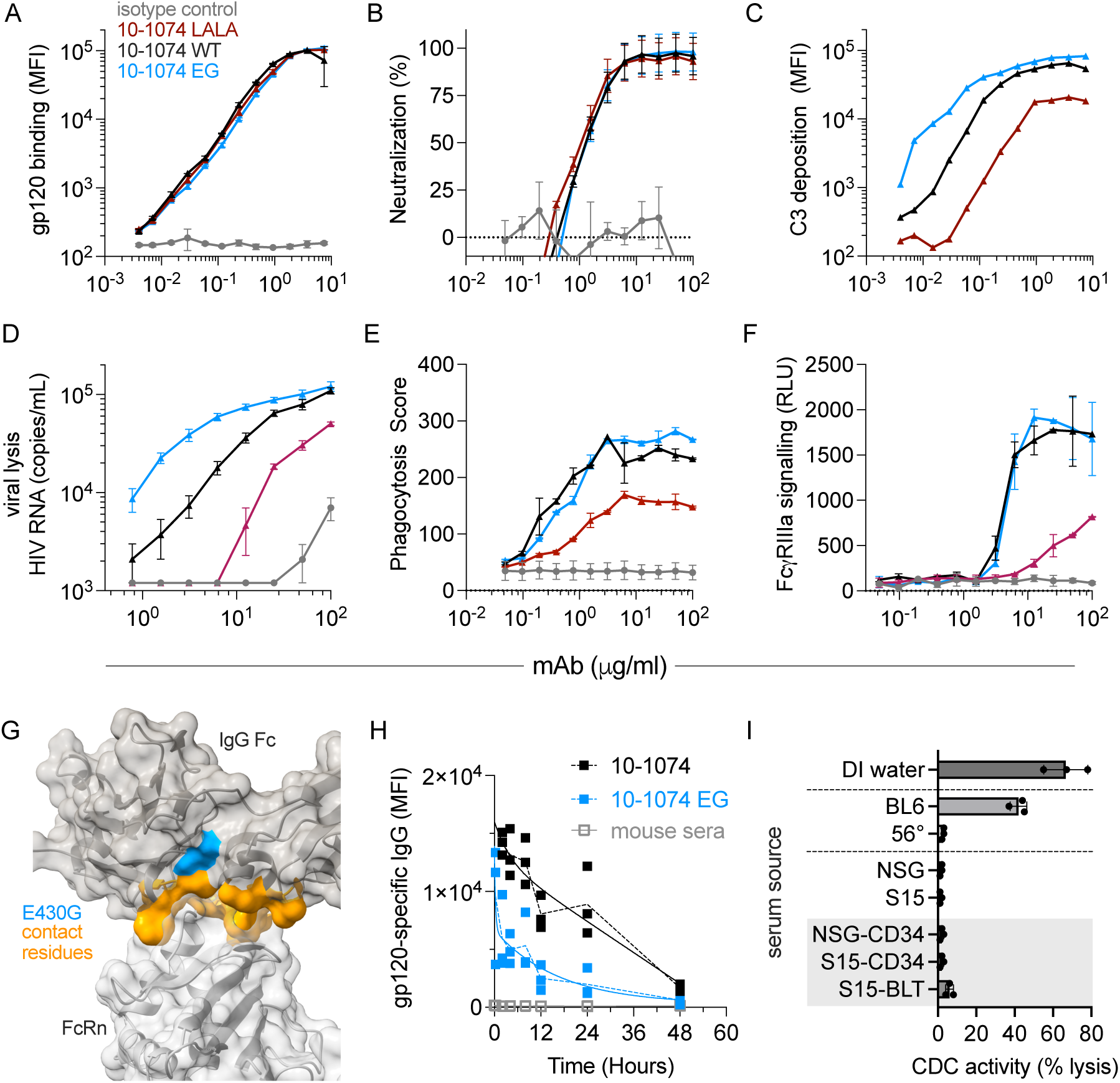
10-1074 Fc variants have different effector functions and pharmacokinetic properties in humanized mice. **A.** Binding of 10-1074 variants and isotype control mAbs to recombinant gp120 in a multiplex binding assay. **B.** Neutralization potency of 10-1074 variants and isotype control mAbs to REJO.c in TZM-BL assay. **C-F.** Effector functions of 10-1074 variants and isotype control mAbs in C3 deposition (**C**), viral lysis in the presence of complement (**D**), phagocytosis (**E**), and ADCC defined as FcγRIIIa signaling (**F**). Error bars indicate standard error of the mean across replicates. **G**. Structural visualization of human IgG Fc (gray) in complex with human FcRn (white) from PDB 7Q15. E430G (blue) and contact residues (orange) are indicated. **H.** Levels of 10-1074 detected in blood over time. **I**. Complement-dependent cytotoxicity (CDC) activity against hemolysin-sensitized sheep red blood cells in serum from indicated controls or mouse strain backgrounds with (gray background) and without humanization. Deionized (DI) water and heat inactivated (56°) mouse serum served as positive and negative controls, respectively.

Though E430G-mutated IgG1 has been reported to exhibit similar clearance as unmodified IgG1 in mice^54^, more rapid clearance has been observed in NHP^39^, consistent with proximity of this position to contact residues involved in pH-dependent binding to the IgG recycling receptor, FcRn, that is responsible for imparting IgG1 with its long half-life (**Figure 2G**). The pharmacokinetic properties of EG and the unmodified IgG1 form of 10-1074 in S15 mice (n = 3 per group) were defined by repeated blood sampling following administration of a 100 μg dose. As was observed in the nonhuman primate model^39^, 10-1074 EG showed more rapid clearance than did 10-1074 IgG1 in S15 mice (**Figure 2H**).

Conveniently, the hemolytic complement (Hc) gene in both NSG and S15 mouse backgrounds is disrupted^55^, resulting in the inability to express C5, a complement cascade protein critical for assembling the membrane attack complex. Terminal complement activation is thus functionally absent in both NSG and S15 mouse strains—thereby providing the opportunity to test whether consistent protection resulted from forms of 10-1074 with variable clearance rates. Lack of complement-dependent cytotoxic (CDC) activity present in serum from NSG and S15 mice, in addition to humanized NSG-CD34, S15-CD34, and S15-BLT mice, was confirmed using hemolysin-sensitized sheep red blood cells (**Figure 2I**).

### Protection from mucosal HIV challenge for 10-1074 IgG1 Fc variants

To define whether similar degrees of protection in the pre-mixing model were offered by forms of 10-1074 with distinct degrees of plasma persistence, two batches of S15-BLT, and one batch of S15-CD34 mice (n_total_ = 65) were challenged intraperitoneally with as escalating dose of transmitted founder REJO.c virus incubated with a 100 μg dose of 10-1074 unmodified IgG1, or EG variant, or 100 μg of CH42 IgG1 isotype control (**Figure 3A**). Infection was monitored over time by evaluation of plasma viremia in blood samples drawn on alternating days given the more rapid progression of infection resulting from systemic challenge.

**Figure 3.**
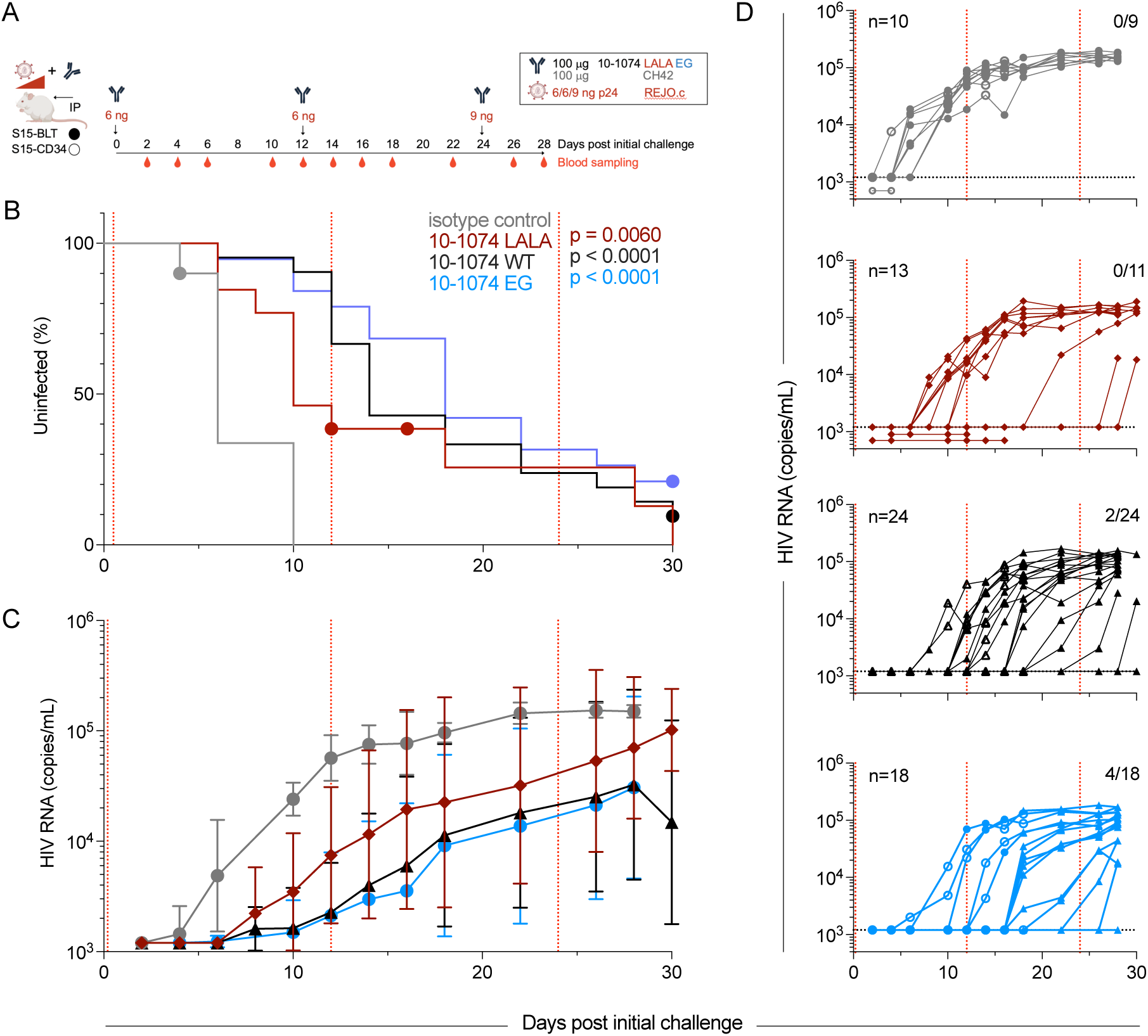
Protection by 10-1074 IgG1 Fc variants can be evaluated despite only moderate in-vitro neutralization and variable pharmacokinetics. **A.** Challenge study design. S15-BLT (filled) or S15-CD34 (hollow) mice were serially challenged with a cocktail of REJO.c virus and mAb intraperitoneally (IP) as illustrated. **B**. Kaplan-Meier curve of infection over time. Statistical significance of 10-1074 treatment compared to isotype control defined by log-rank test. Symbols indicated censored animals. **C-D**. Viral RNA levels over time. Geometric mean and 95% CI of viral RNA over time in each treatment group (**C**) and individual animals (**D**). Dotted horizontal line indicates limit of detection. Dotted vertical lines indicate virus challenge. Animals per group indicated in inset at left, and those uninfected at the end of the experiment in inset at right.

As compared to isotype control, all forms of 10-1074 - unmodified (p < 0.0001), LALA (p = 0.0060), and EG (p < 0.0001) - imparted resistance to infection (**Figure 3B**), extending our observation of antibody-mediated protection from mucosal to systemic challenge routes, and from the CD4 binding site-specific antibody VRC01 to the third hypervariable loop (V3) glycan supersite-specific antibody 10-1074. While time to infection varied, viral loads reached similar levels in infected animals across groups (**Figure 3C**). Despite its inferior pharmacokinetic profile, 10-1074 EG provided a similar degree of protection as unmodified 10-1074 (p = 0.3); two and four of the 24 and 18 animals in the unmodified and EG 10-1074 treatment groups, respectively, remained uninfected at the end of the experiment (**Figure 3D**). Protection conferred by the Fc attenuated 10-1074 LALA variant was inferior when compared to the EG and unmodified antibodies, albeit with limited statistical confidence (p = 0.077). None of 13 animals in the LALA treatment group remained HIV-free, whereas collectively six of 42 animals in the unmodified and EG groups were aviremic following the challenge series.

Additionally, despite differences between the models, both BLT and CD34 humanization mice showed similar degrees of susceptibility to infection and protection mediated by 10-1074 variants. These results stand in stark contrast with results from passive transfer of these forms of 10-1074 and infection following repeated low dose virus challenge in NHP, in which the faster clearance of the EG variant resulted in earlier time to plasma viremia, despite being functionally-enhanced and able to protect at a lower serum concentration^39^.

### Systemic protection of humanized mice from transmitted founder virus by VRC01 depends on IgG subclass

The ability of *in vivo* models to provide value beyond *in vitro* tests relies on their ability to faithfully recapitulate the more complex interactions present between host and pathogen. To this end, both prior NHP and humanized mouse model experiments have shown that beyond direct viral neutralization, antibody-mediated effector functions can contribute to the antiviral effects of monoclonal antibodies *in vivo* (reviewed in^56^). Thus, we next sought to determine the ability of the pre-mixing model to capture these effects and designed experiments around a set of human IgG subclass-switched VRC01 variants.

Consistent with prior reports^57,58^, subclass-switching did not impact antibody binding to HIV-1 envelope protein (**Figure 4A**) or have a major impact on neutralization activity *in vitro* (**Figure 4B**). In contrast, each IgG subclass demonstrated the expected differences in antibody binding profiles to FcγRs as measured by multiplex assay, in which IgG1 and IgG3 exhibit better binding than IgG2 and IgG4 (**Supplemental Figure 2**). Consistent with this interaction profile, IgG1 and IgG3 also exhibited more robust phagocytosis (**Figure 4C**), FcγRIIIa stimulation as a proxy of ADCC activity (**Figure 4D**), and complement-mediated viral lysis (**Figure 4E**). Taken together, these data establish equivalent HIV binding and neutralizing potential but varying Fc-mediated effector functionality across subclasses—demonstrating the suitability of this panel to assess the ability of the pre-mixing model to detect contributions of mechanisms beyond direct viral neutralization.

**Figure 4.**
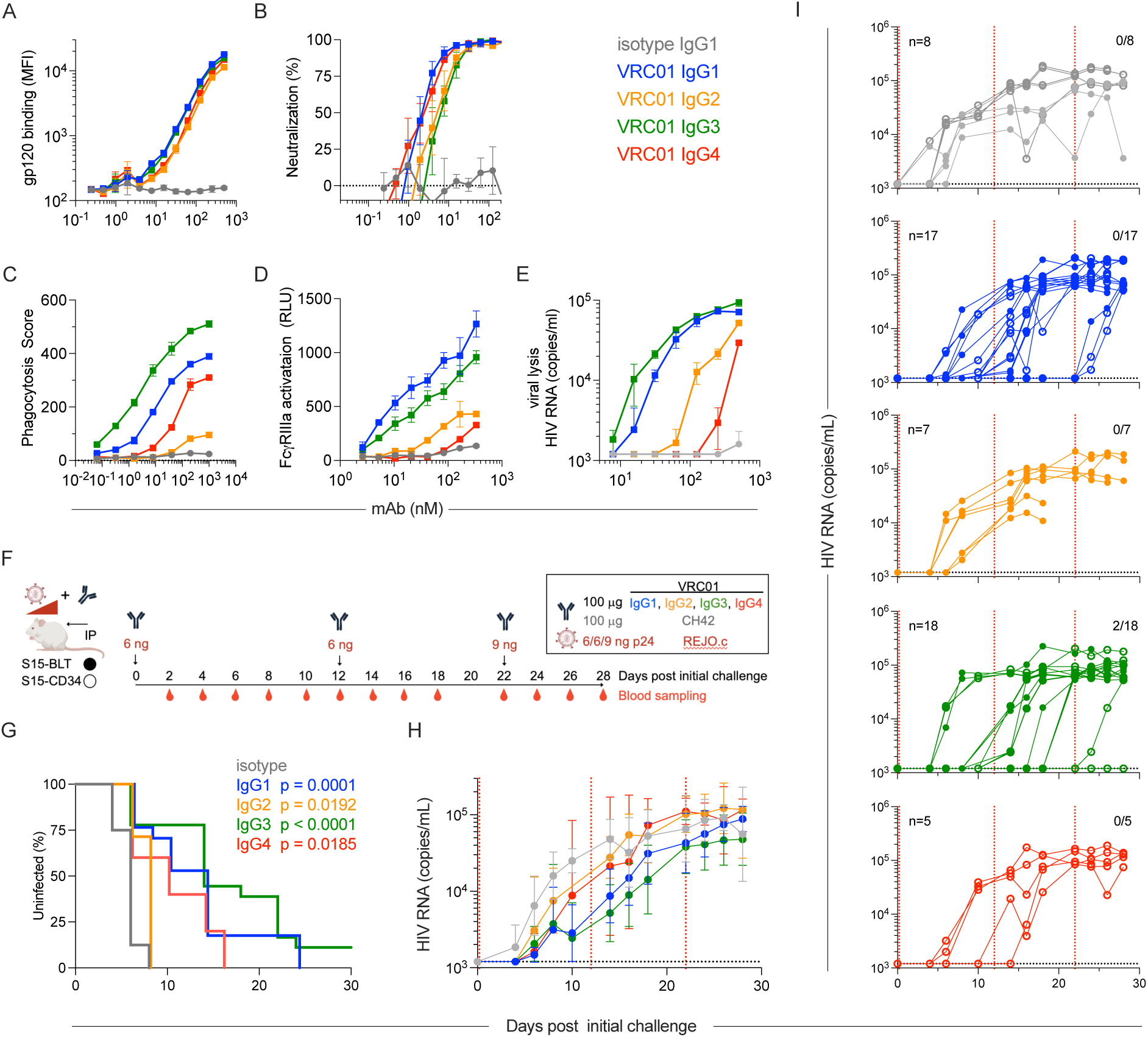
Systemic protection of humanized mice from REJO.c by VRC01 depends on IgG subclass. **A.** Binding of VRC01 IgG subclass variants and IgG1 isotype control mAbs to recombinant gp120. **B.** Neutralization potency of VRC01 IgG subclass variants and IgG1 isotype control mAbs to REJO.c in TZM-BL assay. **C-E.** Effector function of VRC01 subclass variants and isotype control mAbs in (**C**) phagocytosis, (**D**) FcγRIIIa activation, and (**E**) viral lysis in the presence of complement assays. Error bars indicate standard deviation. **F.** Study design. NSG-BLT mice were serially challenged with a cocktail of REJO.c virus and mAb as illustrated. **G**. Kaplan-Meier curve of infection over time. Statistical significance of VRC01 treatment compared to isotype control defined by log-rank test. **H-I**. Viral RNA levels over time. Geometric mean and 95% CI of viral RNA over time in each treatment (**H**) group and (**I**) individual animals. Dotted horizontal line indicates limit of detection. Dotted vertical lines indicate virus challenge. Animals per group indicated in inset at left, and those uninfected at the end of the experiment at right.

One batch of BLT-NSG and another of S15-CD34 humanized mice (n_total_ = 55) underwent repeated intraperitoneal challenge with an escalating dose of the transmitted founder virus REJO.c following pre-incubation with a 100 μg dose of VRC01 in IgG1, IgG2, IgG3, or IgG4 subclasses or with CH42 (IgG1) isotype control (**Figure 4F**). Whereas all animals that received isotype control inoculum were infected by six days post first challenge, all VRC01-treated groups showed delayed plasma viremia (**Figure 4G-H**). Among the subclasses, IgG2 demonstrated only a very modest delay, and like the control group, all animals in this group were also infected following the first challenge (**Figure 4G,I**). In contrast, the other subclasses provided more robust protection in a rank order consistent with their *in vitro* effector function (IgG3 ≥ IgG1 > IgG4). Among these subclasses, IgG1 and IgG3, but not IgG4, exhibited enhanced antiviral activity as compared to IgG2 (p = 0.0022, p = 0.0079, and p = 0.16, respectively by log-rank test). While significant differences in infection risk mediated by IgG3 as compared to IgG1 were not observed (p = 0.16) a greater number of animals were uninfected at the conclusion of the experiment in the IgG3 (2 of 18) than the IgG1 (0 of 17) group (**Figure 4I**). While some comparisons are confounded by group imbalances across batches of humanized mice, infection rates and timing for animals in groups that were well powered in both batches did not suggest systematic differences by batch or humanization strategy. In sum, these data extend previous reports of the impact of subclass switching of VRC07^29^ to its clonal relative VRC01, and to systemic challenge.

### Mucosal protection of humanized mice from transmitted founder virus by VRC01 depends on IgG subclass

Next, because the impact of effector functions may be more easily detected following systemic challenge than in tissue, we sought to define whether subclass-associated differences could also be observed following mucosal challenge with pre-mixed virus and VRC01 inoculum. A series of experiments were conducted to determine whether the subclass-dependent protection afforded by vectored delivery of VRC07^29^, a clonal relative of VRC01, against intravaginal challenge could be extended to challenge with pre-mixed virus and recombinant VRC01 antibody inoculum.

NSG-BLT and S15-BLT mice (n_total_ = 44) were repeatedly challenged intravaginally with 100 μg of VRC01 in IgG1, IgG2, or IgG3 subclasses, or 100 µg or 400 μg of CH42 IgG1 isotype control antibody (in S15-BLT and NSG-BLT batches respectively) when pre-mixed with an escalating dose (3, 9, 27 ng of p24) of transmitted founder REJO.c virus (**Figure 5A**). IgG4 exhibited an intermediate phenotype in the systemic challenge model and so was not tested in the mucosal model. As in the prior intravaginal challenge experiments (**Figure 1**), low dose virus (3 ng p24) did not result in infection of any animals (**Figure 5B**).

**Figure 5.**
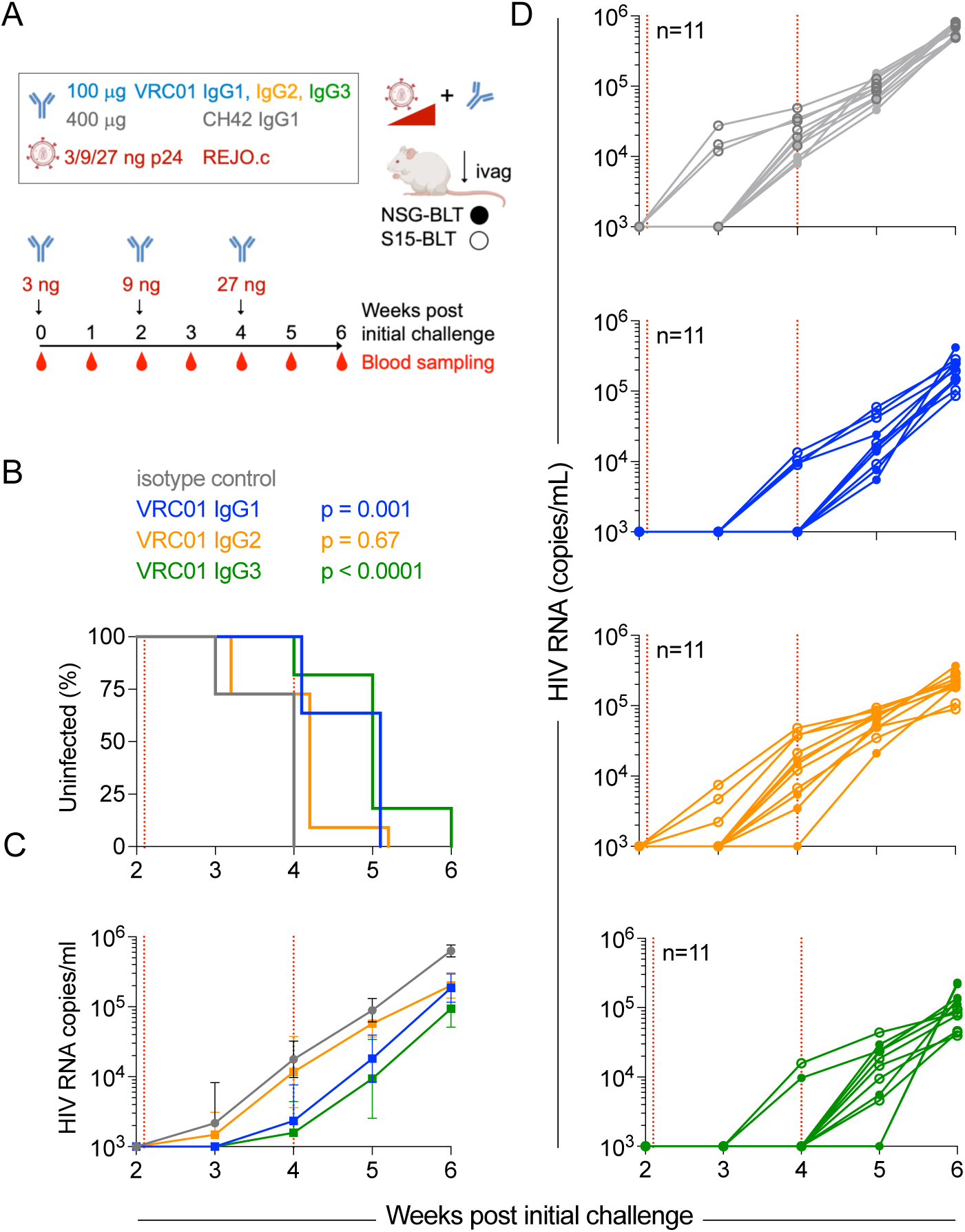
Mucosal protection of humanized mice from REJO.c by VRC01 depends on IgG subclass. **A.** Study design. NSG-BLT (filled symbol) or S15-BLT (hollow symbol) mice were serially challenged intravaginally (ivag) with a cocktail of REJO.c virus and mAb as illustrated. **B**. Kaplan-Meier curve of infection over time. Statistical significance of VRC01 treatment compared to IgG1 isotype control defined by log-rank test. **C-D**. Viral RNA levels over time. Geometric mean and 95% CI of viral RNA over time in (**C**) each treatment group and (**D**) individual animals. Dotted horizontal line indicates limit of detection. Dotted vertical lines indicate virus challenge. Animals per group indicated in inset.

Following the moderate 9 ng virus dose used in the second challenge at week 2, all animals in the isotype control group were infected, as were all but one animal in the IgG2 group (**Figure 5B**). Systemic viremia was observed in approximately 25% of both isotype control and VRC01 IgG2 group animals one week after this challenge. In contrast, both IgG1 (p = 0.001) and IgG3 (p < 0.0001) treatment groups showed significant delays in detection of plasma viremia as compared to isotype control (**Figure 5B,C**). Indeed, IgG3 and IgG1 groups also demonstrated significantly improved protection (p = 0.0006, p = 0.0047 respectively) as compared to IgG2. Though 4/11 mice in the IgG1 group and 2/11 in the IgG3 group were infected at week 4, and 11/11 IgG1 and 10/11 IgG3 group animals were infected at week 5 after the high dose challenge (**Figure 5D**), these differences in time to infection were not significant (p = 0.14). Overall, in combination with the data from systemic challenge experiments, these results demonstrate that the degree of protection afforded by neutralizing antibody is impacted by IgG subclass even when animals are challenged with pre-mixed virus and antibody. The IgG2 form of VRC01 offered inferior protection as compared to IgG1 and IgG3 forms. These results also suggest that IgG3, which demonstrated potentiated phagocytic activity, may provide marginally improved protection as compared to IgG1. Some of the experiments comparing these subclasses were intentionally better powered based on both their greater novelty and potential relevance to the efficacy of the RV144 vaccine regimen^59,60^, as well as the likelihood that differences between these highly effector active subclasses would be smaller than those observed for the more dramatically functionally-compromised IgG2 and IgG4 subclasses. Overall, by demonstrating that antibody effector functions can contribute to the protection afforded by a broadly neutralizing antibody in challenges of pre-mixed antibody and virus administered by systemically and mucosally, these subclass switching experiments establish the model as offering mechanistic insight beyond the *in vitro* neutralizing potential of a given antibody intervention.

### Protection afforded by human serum polyclonal IgG

The reduced quantities of antibody used in the pre-mixing model offer the prospect to more easily conduct experiments to define the protection afforded by passive transfer of polyclonal human antibody pools. To begin to explore this use case, we sourced sera from persons living with HIV (clade B) that exhibited robust *in vitro* neutralization against heterologous clade B pseudoviruses. Among these, IgG purified from sera from donor HIV-008, an anti-retroviral therapy naïve individual with an eight-year history of infection, showed binding to recombinant envelope (**Figure 6A**) and potent neutralization of Tier 2 virus RHPA (**Figure 6B**). Pooled serum IgG purified from seronegative donors (IVIG) was used as a negative control and VRC01 (IgG1) as a positive control. As compared to donor HIV-008, pooled serum IgG purified from seropositive donors (HIVIG) showed similar antigen binding, but considerably poorer neutralization.

**Figure 6.**
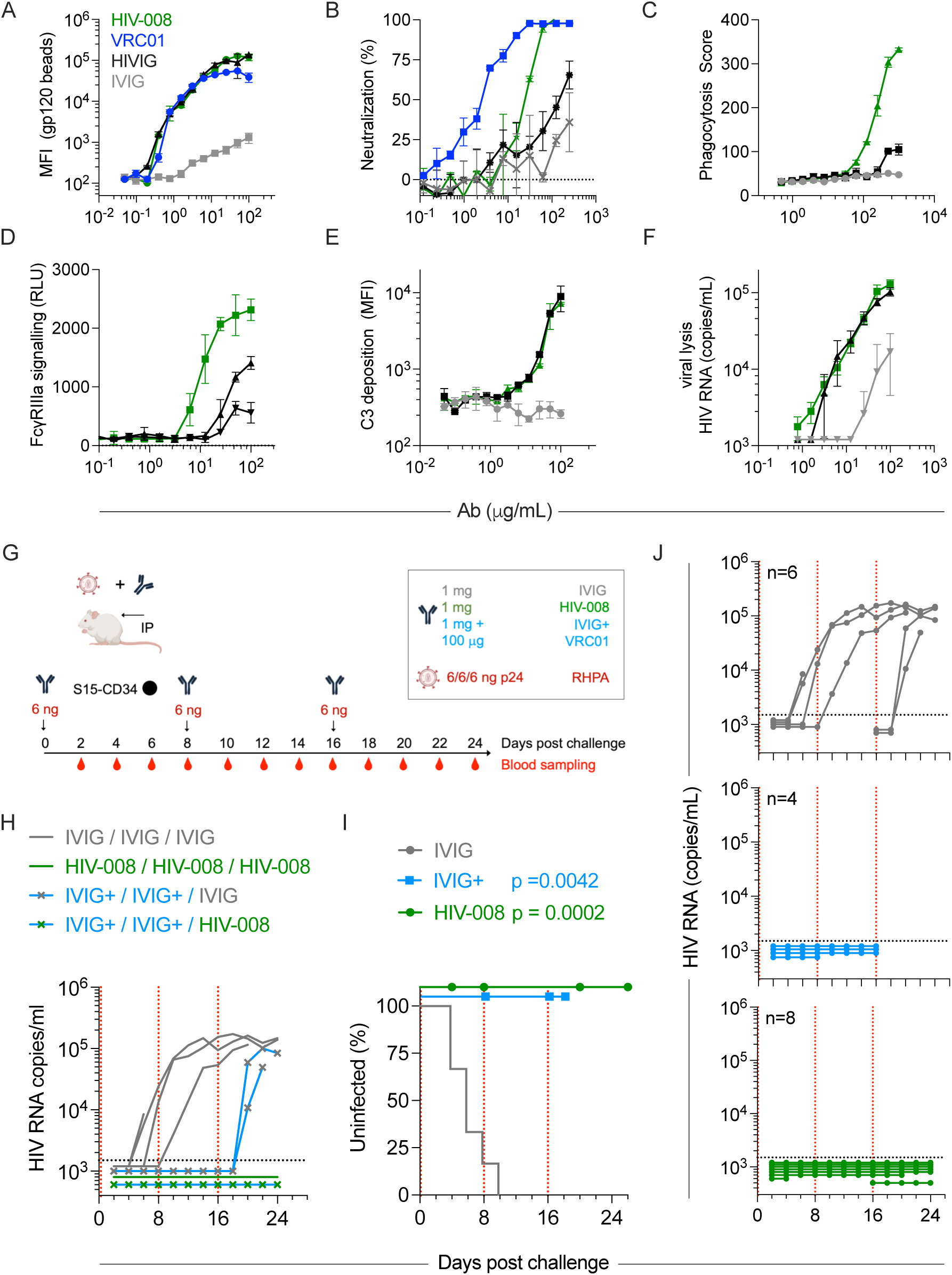
Systemic protection of humanized mice from RHPA by human polyclonal antibodies purified from serum. **A-F**. In vitro antibody activities. **A.** Binding of pooled human serum polyclonal IgG from seronegative donors (IVIG), seropositive donors (HIVIG), from a single seropositive donor (HIV-008), as compared to VRC01 to recombinant gp120 TRO. **B.** Neutralization potency against RHPA in TZM-BL assay. **C-F.** Effector function of pAbs in phagocytosis (**C**), ADCC defined by FcγRIIIa activation (**D**), complement deposition (**E**), and complement-dependent viral lysis (**F**) assays. Error bars indicate standard deviation. **G.** Study design. One batch of S15-CD34 mice was serially challenged intraperitoneally (IP) with a cocktail of RHPA virus and pAb as illustrated. Animals in the IVIG plus VRC01 group were randomized to either IVIG or HIV-008 after the second challenge. **H-J**. Detection of viral RNA over time by (**H**) treatment group and (**J**) per intervention. Dotted horizontal line indicates limit of detection. Animals per group are indicated in inset. **I**. Kaplan-Meier curve of infection over time. Results from the third challenge of animals re-randomized from the IVIG plus VRC01 group after the second challenge were included with newly assigned groups at challenge one. Statistical significance of IVIG plus VRC01 and HIV-008 treatment compared to IVIG defined by log-rank test. Censored animals indicated by symbols. Dotted vertical lines indicate times of virus challenge.

Assessment of *in vitro* effector function of polyclonal IgG purified from donor HIV-008 showed greater phagocytosis (**Figure 6C**) and ADCC (**Figure 6D**) activity, but not C3 deposition (**Figure 6E**) or complement-mediated lysis (**Figure 6F**) compared to HIVIG. With the exception of complement-mediated lysis, which was orders of magnitude reduced in activity, IVIG generally showed undetectable effector function.

The ability of these polyclonal pools to protect from systemic challenge was then evaluated in S15-CD34 mice (n_total_ = 18). Plasma viremia was assessed in mice challenged intraperitoneally with a series of up to three repeated moderate (6 ng of p24) doses of RHPA virus pre-mixed with either purified polyclonal IgG from donor HIV-008 at a dose of 1 mg (n = 6), 1 mg of IVIG (n = 4), or 1 mg of IVIG plus 100 μg of VRC01 (n = 4) (**Figure 6G**). After the first challenge, 3/4 IVIG-treated mice were infected, whereas no mice treated with IVIG plus VRC01 or HIV-008 IgG exhibited detectable plasma viremia (**Figure 6H**). After the second challenge, the remaining IVIG-treated mouse was infected, while no animals treated with HIV-008 or with IVIG plus VRC01 exhibited viremia. Given this striking result and the group imbalance in the initial design, for the third challenge, the three surviving positive control group mice were randomized back out to IVIG and HIV-008 groups to improve power. Both mice randomized to IVIG were infected following the third challenge; the lone animal switched to HIV-008 remained uninfected (**Figure 6I**). In survival analysis in which re-randomized mice were considered as providing insight into both a series of two challenges in the context of IVIG plus VRC01 as well as a single challenge in their randomized treatment arm, both the positive control group (IVIG plus VRC01) and the HIV-008 sera groups showed significant protection (**Figure 6J**, p = 0.0042 and p = 0.0002, respectively). These data demonstrate that polyclonal human serum IgG with robust neutralization and effector function can protect humanized mice from virus challenge.

## Discussion

Identification of both virological and immunological correlates of risk (CoR) in case-control study designs suggest that some individuals have been protected from some HIV-1 viruses in recent field efficacy trials of both active and passive immunization. Yet, substantial improvements will be needed for these observations to translate into approved vaccines and antibody prophylactics. The ability of these footholds to support more rapid development of promising interventions to prevent HIV infection with greater efficacy depends in part on facile and reliable preclinical models. With this goal in mind, we tested the ability of simple adaptations of *in vivo* antibody protection and virus challenge strategies in the humanized mouse model to reduce biological and technical variability while maintaining the ability to evaluate mechanisms of antibody-mediated protection from HIV.

Firstly, escalating virus challenge dose over a series of repeated mucosal challenges supported efficient modeling of relationships between antibody and virus dose. Over doses of each that varied by approximately an order of magnitude, dose-dependent protection profiles were consistently observed. Using this strategy, dedicated batches of mice were not needed to establish either *in vivo* midpoint infectious dose (ID_50_) values for each virus or midpoint protective dose values for antibody. The efficacy of control and test interventions could be evaluated at the same time, and these benchmarks were defined using approximately one-tenth the amount of recombinant antibody as is typically administered in a one-time administration to a single NHP.

Secondly, pre-mixing antibody and virus supported meaningful comparative testing of antibodies with divergent PK properties. Here, this limitation was modeled with 10-1074 EG, an Fc mutated form of 10-1074 that exhibited rapid clearance in both mice and NHP and had failed to protect NHP from serial challenges as durably as 10-1074 lacking Fc modification^39^. Despite more rapid clearance when administered systemically in mice, Fc modified 10-1074 EG showed equivalent protection against systemic challenge as unmodified 10-1074 when virus and antibody were pre-mixed. We posit that by providing a known and consistent quantity of antibody at its site of action, pre-mixing reduces the biological variability within treatment groups that is associated with individual-specific antibody biodistribution and clearance rates. This experiment also showed that the Fc-compromised 10-1074 LALA exhibited marginally poorer protection than fully functional counterparts.

Relative to more standard humanized mouse experimental designs, pre-mixing also reduced the amount of monoclonal antibody needed by an order of magnitude - from 1 mg to 0.1 mg per animal per challenge. Decreased material requirements made it feasible to test the efficacy of some eight different monoclonal antibodies—most in both mucosal and systemic challenge settings—with quantities easily prepared in an academic lab setting. Lowering this barrier to experimental design permitted comparison of a subclass-switched set of VRC01 variants. This testing extended the previous report of the impact of subclass switching of VRC07^29^, which was found to depend on Fc-domain effector functions for optimal protection, to include VRC01. Discrimination between subclasses was made with relatively fewer animals, as group comparisons in the VRC07 study were confounded by different levels of antibody that resulted from adeno-associated virus-vectored delivery of antibody. Here, in both systemic and mucosal challenge settings, IgG subclasses with compromised effector function provided compromised ability to protect from infection. Like that for the 10-1074 variants, this differentiation demonstrates that not only are these activities important to protection afforded by VRC01, but that the pre-mixing model captures a broad set of antiviral effects, such as innate immune effector functions that vary across human IgG subclasses in ways that are relevant to reducing risk of HIV infection.

Importantly, the humanized mouse model proved responsive to both systemic (intraperitoneal) and mucosal (intravaginal) challenge routes, highlighting its versatility in mimicking different routes of HIV exposure and offering a biologically relevant context to evaluate mucosal and systemic immunity. These findings support this experimental approach in the humanized mouse model as a valuable resource for understanding antibody activities beyond neutralization and underscore its potential to guide Fc engineering strategies in HIV treatment and prevention. The model’s responsiveness to graded variations in antibody Fc characteristics provides a unique avenue to preclinically evaluate and refine Fc-engineered therapeutics, enhancing our ability to optimize immune engagement and antiviral efficacy and contribute to the rational design of next-generation antibody therapies.

Perhaps most promisingly, these results supported the feasibility and potential value of evaluating protection afforded by polyclonal human serum IgG. Aiming for similarly potent *in vitro* neutralizing activity as was achieved by protective monoclonal antibodies, we calculated that 1 mg of serum IgG from a well-characterized seropositive donor would provide similar neutralizing activity as a protective dose of VRC01. As this quantity of serum IgG can typically be collected from 0.1 mL of serum, just a few mL of blood were sufficient to support testing eight animals against a series of three repeated challenges.

To this end, human HIV-1 vaccine efficacy trials have identified multiple humoral correlates of reduced infection risk^60–65^. While some of these correlates have been at least somewhat bolstered by NHP studies^66–74^, direct mechanistic validation of humoral correlates observed in humans by conducting passive transfer studies has not been feasible. Taking these polyclonal humoral responses back into preclinical settings for validation has been challenged by technical factors as well as the low power anticipated to result from the only moderate magnitude of effects observed in humans. As a result, with the exception of protection mediated by VRC01, which has a rich evidence base in preclinical models^75^ and clinical support from the AMP trials^5^, no human humoral correlate of protection (CoP) is supported by better than associational evidence. Direct mechanistic testing of the humoral correlates of reduced risk observed in human trials has critical importance to defining whether or not humoral CoP such as IgG3^59–61,65^, ADCC^62^, or variable loop binding^62–64^ responses hold independent or added value to bnAb-based vaccine and Ab prophylaxis strategies. Our observation that humanized mice can be protected from infection by polyclonal human serum antibodies provides not only a practical demonstration of efficient efficacy testing, but the first evidence that naturally-derived and expressed pools of human antibodies can prevent HIV-1 infection *in vivo*. Following this work with tests of vaccine-elicited serum antibodies as compared to placebo controls, and with sensitive and non-sensitive viral strains^8,9^ will be important to further demonstrate both the practical utility and the scientific value that can be realized by these approaches to simplifying and de-risking *in vivo* experiments.

Nonetheless, further challenges remain and further optimizations are possible. While challenge with serendipitously^76^ or intentionally mixed pools of virus has the potential to yield insight into efficacy against more than a single virus in a given experiment, such approaches are only now beginning to be developed. The presence and functional capacity of various effector cells in mucosal tissue and comparability to humans in these models are incompletely characterized. Neutrophils are not adequately reconstituted in the humanized models used. Additionally, there are limitations to the work presented here. Different numbers of animals were used across groups, yielding different power to detect differences. Some batches of mice included censored animals that succumbed to apparent graft versus host disease (GVHD) before completion of the observation period. Some experiments terminated before viremia reached steady state in all animals, precluding comparison of effects on this outcome. Viral load data were not successfully collected from all animals at all draw timepoints; and in some cases, days of draw were imperfectly consistent between batches of animals. Though time to infection supported reasonable inferences as to which challenge resulted in systemic viremia, these inferences could not be proven. While humanization profiles cleared a minimum threshold, detailed enumeration of various target and effector cell populations, and their variability from animal to animal or batch to batch was not defined and therefore could not be related to infection outcomes. Some experimental design decisions were made based on observations of GVHD, loss of mice, or outcome information from the prior challenge. Lastly, a single human polyclonal serum antibody sample was tested in a single batch of animals.

Limitations notwithstanding, insights derived from more resource-efficient *in vivo* models are desperately needed to accelerate the pace of progress in developing humoral interventions to prevent HIV-1 infection. In humans, the limited number of both active as well as passive vaccine efficacy trials conducted to date, open questions about analysis and interpretation of their outcomes^77^, and the unclear mechanistic relevance of humoral CoP stand as impediments to progress. In theory, humanized mouse models offer an economical and practically accessible means to model protection from HIV afforded by vaccine-elicited or recombinant antibodies *in vivo*. However, all *in vivo* models present high-stakes decision points that, if not chosen appropriately, pose considerable risk to the ability of studies to address their research objectives. Collectively, this work demonstrates methodological approaches that promise to lower barriers to further exploration of antiviral antibody mechanism of action and protection from HIV-1 afforded by human polyclonal antibody responses induced by even vaccination.

## Materials and Methods

### Antibody and virus preparations

#### Expression and purification of HIV-specific and control antibodies

VRC-HIVMAB060-00-AB (VRC01) IgG1 was provided by the NIH Vaccine Research Center, and was produced as described in the protocols for the HVTN 703 and 704 clinical trials^5^. Briefly, sequences encoding for variable region of VRC01 light and heavy chain were codon optimized and stably transfected into Chinese hamster ovarian (CHO) cells. This engineered CHO cell line, was used to express VRC01 under cGMP standards. Subclass-switched variants of VRC01^78^, including IgG2, IgG3s with a substitution made to increase stability (N392K)^79^, and IgG4a1, Fc engineered forms of 10-1074^39^ including IgG1, 10-1074 LALA [L234A/L235A], and 10-1074 EG [E430G], and Herpes Simplex Virus glycoprotein D-specific IgG1 isotype control mAb CH42^80^ were produced using either Expi293 or ExpiCHO expression systems (Thermo). Briefly, heavy and light chain plasmids were transiently co-transfected by polyethyleneimine or according to the ExpiCHO transfection kit (Thermo Fisher) protocol. Culture supernatants were collected at 10 days post-transfection, purified by gravity using MabSelect protein A chromatography resin (GE Healthcare), except VRC01 IgG3 which was purified using MabSelect protein G chromatography resin (GE Healthcare). All procedures were performed in a sterile biosafety cabinet and antibodies were buffer exchanged into PBS using 10kDa molecular weight cut off columns (Millipore). Antibody preps were tested for endotoxin using the ToxinSensor kit (Genscript). If necessary, endotoxins were removed using Pierce High Capacity Endotoxin Removal Spin Columns (Thermo Scientific). All final antibody preparations were formulated in PBS and had endotoxin levels below 0.1EU/ml.

#### Polyclonal serum IgG sourcing and evaluation

Blood samples from people living with HIV that were not on anti-retroviral therapy were collected from voluntary donors who provided written informed consent at Brigham & Women’s Hospital (Boston, MA) under IRB (2006P001197). Serum samples were obtained following standard blood processing procedures and aliquots stored at −80’ C until use. Donor serum samples were screened in TZM.bl neutralization assays against a standard reference panel of clade B HIV-1 Env pseudoviruses^81^ to characterize neutralizing activity. Donor HIV-008, selected on the basis of broad and potent neutralizing activity that included the REJO strain, was a 45 year-old female participant that had been diagnosed with HIV 8 years prior to blood sample collection, and had never been on antiretroviral therapy.

#### HIV production and quantification

Plasmids encoding for infectious molecular clones (IMC) of HIVpREJO.c [Balazs Lab; BEI HRP-11746] or HIVpRHPA [Seaman Lab; BEI HRP-11744] were replicated in E.coli [NEB Stable] and sequence confirmed following maxiprep. Infection and replication competent HIV were generated by transiently transfecting HEK293T cells with IMC plasmid. Briefly, HEK293T cells were cultured in T150 flasks to ∼80% confluency and 20ml of media [DMEM + 10% FBS]. 150µl of IMC plasmid (2 mg/mL) was added to 1,350 µL of Opti-MEM (Invitrogen), and mixed with 1,500µl of 1:10 diluted Lipofectamine 2000. This mixture was incubated for 5 minutes at room temperature, and then 2.5mL were added to the HEK293T cell culture flask. Cells were incubated for 48hours. Cell supernatant was collected, centrifuged at 300 x g for 10 minutes to separate cells and debris, filtered through a 0.45 µm filter, and frozen in 1ml aliquots. Each preparation of viral stock was characterized for p24 quantity using a p24 ELISA assay [Leidos Biomedical Research]. Viral stocks were also assayed for infectivity by titrating infection in-vitro on TZM-bl cells [BEI Resources] to calculate a 50% tissue culture infective dose (TCID_50_) using the Spearman-Karber formula.

### Humanized mouse challenge experiments

#### Preparation of humanized mice

Immunodeficient NOD/SCID IL2Rgammanull (NSG) mice and NSG-SGM3-IL15 (S15) mice were obtained from The Jackson Laboratory and humanized by the Animal Cancer Models resource at Dartmouth Cancer Center. Briefly, 6- to 8-week-old female NSG or S15 mice were irradiated with 5 Gy γ-radiation and allowed to recover for 4 hours. Following recovery, irradiated mice were transplanted with human fetal liver and thymus tissue under the kidney capsule and injected intravenously with 130,000 – 200,000 CD34+ cells isolated from autologous fetal liver^82^. The resulting BLT mice were rested for 11-12 weeks after surgery to allow for recovery and engraftment. Human CD34+ cell engrafted NSG-CD34 or S15-CD34 mice were generated be adopting a previously reported protocol.^83^ Briefly, 24-72 hour old pups were irradiated with 1 Gy γ-radiation and allowed to recover for 4 hours. Following recovery, 100,000 human CD34+ cells (Stemcell) were administered to the pups by intra-hepatic injection in a 50 µL volume. Mice were rested 11-12 weeks after cell administration to allow for engraftment. All mice were maintained on Septra water and housed in BSL-2 facilities post-surgery. To confirm successful humanization, at this time blood samples were taken from mice by retro-orbital bleeding and centrifuged at 1100 x g for 5 min at room temperature to separate plasma from cell pellets.

Plasma was removed and frozen at −80°C for subsequent analysis as negative serum control. Cell pellets were resuspended in 1 ml of 1× red blood cell (RBC) lysis buffer (BioLegend) and incubated on ice for 10 min. After RBC lysis, each sample was pelleted at 1100 x g in a centrifuge for 5 min at room temperature and then stained with 100 µL of an antibody cocktail containing 1:100 diluted anti-mouse CD45-APC/Fire^TM^ 810 (Biologend, clone 30-F11), anti-human CD45-pacific blue (Biologend, clone HI30), anti-human CD4-AF488 (Biologend, clone RPA-T4), and anti-human CD8-PE (Biologend, clone SK1) in PBSA (PBS supplemented with 1% BSA) for 45 min on ice. Samples were washed by pelleting at 1100 x g and resuspending into 200 µL of PBSA and then analysed on a Novocyte Advanteon flow cytometer. Mice with at least a 20% huCD45+ population were considered sufficiently reconstituted and included in the experiments (**Supplemental Figure 1**). Animal experiments were done with approval from the Institutional Animal Care and Use Committee of Dartmouth College and conducted in accordance with the regulations of the American Association for the Accreditation of Laboratory Animal Care.

#### Viral challenge

Frozen aliquots of HIVpREJO and HIVpRHPA virus were thawed in a 37°C water bath. Desired amounts of virus and antibody were then combined in microcentrifuge tubes and incubated at 37°C for 1 hour. Vaginal challenges were performed by placing isoflurane-anesthetized mice in a supine position and followed by a shallow insertion of p200 pipette to deliver 50 µL of antibody-virus mix into the vaginal vault. Mice were maintained in a supine position with their posterior raised for 5 min to prevent loss of the virus. For intraperitoneal (IP) challenges, isoflurane-anesthetized mice were injected with 100 µL of the antibody-virus mixture using a 28-guage syringe. Retro-orbital blood draws were performed periodically as described above to permit assessment of systemic viremia.

#### Viral load quantification by qPCR

Viral RNA was extracted from mouse samples using the QIAamp Viral RNA Mini Kit (Qiagen). Each RNA sample was treated with 2 U of Turbo DNase (Invitrogen) for 30 min at 37°C, followed by 15 min at 75°C for heat inactivation. Samples were then used to prepare cDNA using QuantiTect Rev. Transcription Kit (Qiagen). These samples were then used in a 20 µL qPCR reaction with SsoAdvanced Universal SYBR® Green Supermix (Biorad) and primers designed to target the Pol gene of HIVREJO.c (5′-CAATGGCCCCAATTTCATCA and 3′-GAATGCCGAATTCCTGCTTGA). Samples were run in triplicate on a Biorad 96-cfm Real-Time PCR system (Biorad) with the following cycle conditions: 95°C for 3 min, followed by 60 cycles of 95°C for 3 s and 70°C for 30 s. Virus titre was determined by comparison with a standard curve generated using RNA extracted from a serially diluted mixture of commercially-titered viral stock (BEI Resources) and undiluted mouse serum. The limit of detection was 1200 HIV copies/mL.

### *In vitro* antibody characterization

#### Multiplex antigen binding assay

Binding of antibodies to HIV envelope protein was assessed by multiplex assay as previously described^84^. Briefly, TRO HIV gp120 (Immune Technology) and HSV-2 gD^2^ [Cohen Lab] were covalently coupled to coded MagPlex superparamagnetic carboxylated magnetic microparticles (Luminex) are previously described^3^. Antigen coated beads were washed prior to incubation with antibody prepared as two-fold dilutions starting at 500 nM in assay buffer (1× PBS, 0.1% BSA, 0.05% Tween 20) for 2 hr at room temperature (RT) with shaking. Plates were washed five times on an automated plate washer (BioTek) and subsequently detected with 0.7 μg/mL phycoerythrin (PE)-conjugated goat anti-human IgG Fc (Southern Biotech) for 1 hr at RT with shaking. Following five wash cycles, beads were resuspended in xMAP sheath fluid (Luminex), and sample median fluorescence intensities (MFIs) were collected using the Flexmap 3D system (Luminex).

#### Neutralization

*In-vitro* neutralization assays were performed as described^29^. Briefly, 96-well plates were prepared with 10,000 TZMbl cells per well and allowed to incubate for 24 hours. After this incubation, the supernatant was removed and replaced with 100µl/well of fresh culture media supplemented with diethylaminoethyl dextran (112.5 µg/mL) to facilitate viral infection. 200 TCID_50_ of HIVpREJO or HIVpRHPA virus was combined with two-fold serial dilutions of antibody in culture media (DMEM + 10% FBS), starting at either 250nM (mAbs) or 250 µg/ml (pAbs), and incubated at 37°C for 1 hour. 50 µl of the virus-antibody mixtures were then added to wells in the previously prepared 96-well assay plate and incubated for 48 hours at 37°C. Post incubation, 150ul of room temperature SteadyGlo reagent (Promega) was added to each well of the assay plate and allowed to incubate for 20 minutes. 200 µl from each well was transferred to a white opaque 96-well plate and luciferase activity was measured on a SpectraMax L microplate reader (Molecular Devices). Background luciferase signal from negative control cells (cells only) was subtracted from all samples. Neutralization activity was calculated as a percentage of the difference between virus control wells (cells with virus only) and negative control wells (cells only).

#### Phagocytosis

Antibody-dependent phagocytic activity was determined by quantification of antigen-conjugated fluorescent beads by the monocytic human THP-1 cell line^85^. Briefly, fluorescent polystyrene 1.0 µm beads (Life Technologies) were conjugated with TRO HIV gp120 (Immune Technology) protein via crosslinking carboxylic bead surface with primary amines. Following washing, 10^5^ beads per well were incubated in media [RPMI1640 (Fisher) + 10% FBS (Biowest] with antibody across a two-fold dilution series starting at 500 nM. THP-1 cells (ATCC) at a 1:10 cell to beads ratio and incubated at 37°C under 5% CO_2_ for 4 hours. Cells were washed in cold PBS, fixed with 4% paraformaldehyde, washed, and analysed on a flow cytometer (Advanteon Novocyte). Flow cytometry analysis included gating on the cell population in the FSC vs SSC plot, followed by identification of the proportion of cells that were bead+ and the median fluorescent intensities (MFI) of this population (**Supplemental Figure 3**). A phagocytosis score was calculated by multiplying the MFI value by the percent bead^+^.

#### Antibody-dependent cellular cytotoxicity

An FcgRIIIA stimulation reported cell assay was used to model ADCC activity. Assay plates were prepared by coating high-binding flat bottom 96-well plates (VWR) with 50 µL per well of 1µg/mL TRO gp120 (BEI resource) in PBS and incubated overnight at 4°C. Assay plates were washed with PBS and then blocked using PBS + 2.5% bovine serum albumin (Sigma) for 1 hour at room temperature. Assay plates were washed with PBS and 20 µL of titrated antibody dilutions were added to each well. A 20 µL volume of PBS was used as a negative control. Assay plates with antibodies were allowed to incubate for 20 min. Jurkat-Lucia NFAT-CD16 cells (Invivogen) were cultured in growth medium - IMDM media supplemented with 10% fetal bovine serum, 1 mM sodium pyruvate, 1x non-essential amino acids, 100 µg/mL Normocin, 100 µg/mL Zeocin, 10 µg/mL Blasticidin. Cells were pelleted by centrifugation at 300 x g for 7 minutes and resuspended in IMDM + 10% fetal bovine serum, 1 mM sodium pyruvate, and 1x non-essential amino acids. A 180 µL volume containing 100,000 cells was added to each well. Assay plates were incubated for 24 hours at 37°C with 5% CO_2_. Following incubation, 25 µL from each well was mixed with 75 µL of Quanti-Luc luciferase substrate (Invivogen) and luciferase signal was measured on a plate reader and analysed using Prism.

#### Antibody-dependent complement deposition

HIV gp120-conjugated Magplex assay beads were prepared as described above and incubated with antibodies titrated from 500 nM to 0.48 nM, for 2hr at RT with constant agitation. Complement active guinea pig serum [Fisher Scientific] was diluted 1:100 in GVB^++^(Sigma) and added to the assay plate and incubated for 30 mins at 37°C. The assay plate was washed with cold PBS to stop the complement reaction and beads then incubated with biotinylated anti-guinea pig C3d antibody (Thermo) tetramerized by mixing 4:1 with streptavidin-PE 20 min at RT before use. Tetramerized anti-C3d-PE was added to assay plate at 0.7 µg/mL and incubated for 1 hr at RT with shaking. Following another wash cycle, beads were resuspended in xMAP sheath fluid (Luminex), and sample median fluorescence intensities (MFIs) were collected using the Flexmap 3D system (Luminex).

#### Antibody-mediated complement-dependent viral lysis

HIV viral particles were packaged as described above. HIV was quick thawed in a 37°C water bath and then diluted in GVB^++^(Sigma) to 100,000 copies/ml of HIV RNA. A 25 µL volume of prepared HIV was then added to each well in a 96-well PCR plate (Biorad). Antibodies were titrated in GVB^++^ from 1000 nM to 0.96 nM and 25 µL of diluted antibodies were added to the PCR plate containing HIV. No HIV and GVB^++^ were used as negative controls while Triton X-100 (Sigma) was used as a positive control. Antibody dilutions and HIV were incubated for 20 min at room temperature. Complement active guinea pig serum (Cedarlane) was diluted 1:100 in GVB^++^ and 50 µL was added to each well in the PCR plate and incubated for 4 hours at 37°C. A 50 µL volume of supernatant from each well was used to prepare cDNA using the QuantiTect Reverse Transcription Kit (Qiagen). These samples were then assayed for HIV copies via targeted qPCR and quantified based on a standard curve as described below.

#### Complement-mediated cell lysis assay

Sheep RBC lysis assay was performed as previously described^86^. Briefly, 1.0 mL of Sheep red blood cells (RBCs) (Innovative Research, ISHRBC100P15ML) were washed three times by suspension in 3.0 ml of GVB++ (Complement Technology, B100) and centrifugation at 900 x g for 5 min. RBCs were then resuspended in 10.0 mL of GVB++ to make a stock solution of 10% sheep RBCs. Next, 10.0 ml of GVB++ containing 1:50 diluted Hemolysin (Complement Technology, Hemolysin) was added to the 10% sheep RBC solution, and this mixture was incubated at 30°C for 30 min with continuous end-over-end mixing. This solution of hemolysin-sensitized RBCs was then stored at 4°C with continuous mixing till the assay was performed. The assay plate was prepared by adding 50 µL of hemolysin-treated sheep RBCs to each well of a 96-well V bottom plate (USA Scientific, 1833-9600). Serum from various mice strains were diluted 1:100 in GVB++, and 50 µl was added to separate wells. Cells treated with hemolysin in advance of two −80°C freeze-thaw cycles and vortexing were used as a positive control for to define 100% lysis while a 50 µL volume of filtered DI water served as an additional positive control and 50 µL of GVB++ alone and 50 µL of heat inactivated serum from BL6 mice were used as negative controls. The prepared assay plate was then incubated at 30°C for 30 min, followed by centrifugation at 1500 x g for 5 min. A 50 µL volume from each sample was then mixed with 50 µL filtered DI water in a 96-well white clear bottom plate (Costar, 3610) and absorbance was measured at 540 nm (SpectraMax Paradigm plate reader, Molecular Devices). Background absorbance was subtracted from all samples and then complement-mediated RBC lysis was then calculated as a percentage of total lysis as defined by the freeze-thawed sample.

### Antibody pharmacokinetics

A 100 µg quantity of either unmodified wild type (WT) or EG mutant 10-1074 antibody was administered to mice via tail vein injection in a volume of 50 µL. Longitudinal blood samples were collected via retro orbital bleeds. Blood samples (50 µL) were spun at 1100 x g for 10 minutes to pellet cells. Serum was collected and frozen at −80. When all serum samples had been collected, samples were thawed and diluted 1:250 and 10-1074 antibody levels were assessed by multiplex assay as described above.

### Structure Visualization

Sequences and coordinates for human IgG1 in complex with FcRn were retrieved from the Protein Data Bank (PDB) entry 7Q15^87^. Contact residues were determined from the combined model using buried solvent-accessible surface area ≥ 15 Å^2^. Structures were visualized using ChimeraX version 1.9^88^.

### Statistical analysis

Data visualization and statistical analysis were performed using Prism (GraphPad Software). Mice were designated as infected when plasma viral load was detected at higher than minimum limit of detection. Log rank tests were used to assess statistical significance.

## Data availability

Study data are openly available at [link to be inserted].

## Acknowledgements

Irradiation, Imaging, Microscopy and Animal Cancer Models Shared Resource are supported by Dartmouth Cancer Center support grant P30CA023108 from the National Cancer Institute. This work was also supported in part by the Gates Foundation INV036842 and administrative supplement to the National Institutes of Health NIAID P01AI162242. The authors have no conflicting financial interests.

