## Supplemental Materials for "Effects and mechanisms of monoclonal and polyclonal human antibodies in protection of humanized mice from HIV-1 challenge"

**Supplemental Figures and Legends**

| **Supplementary Figures** | |
|---|---|
| Supplemental Figure 1 | Method for assaying reconstitution of human immune cell population and sorting animals for experiments |
| Supplemental Figure 2 | VRC01 IgG subclasses binding to human FcγR |
| Supplemental Figure 3 | Gating strategy for assessing antibody-dependent cellular phagocytosis (ADCP) via THP-1 cell assay |

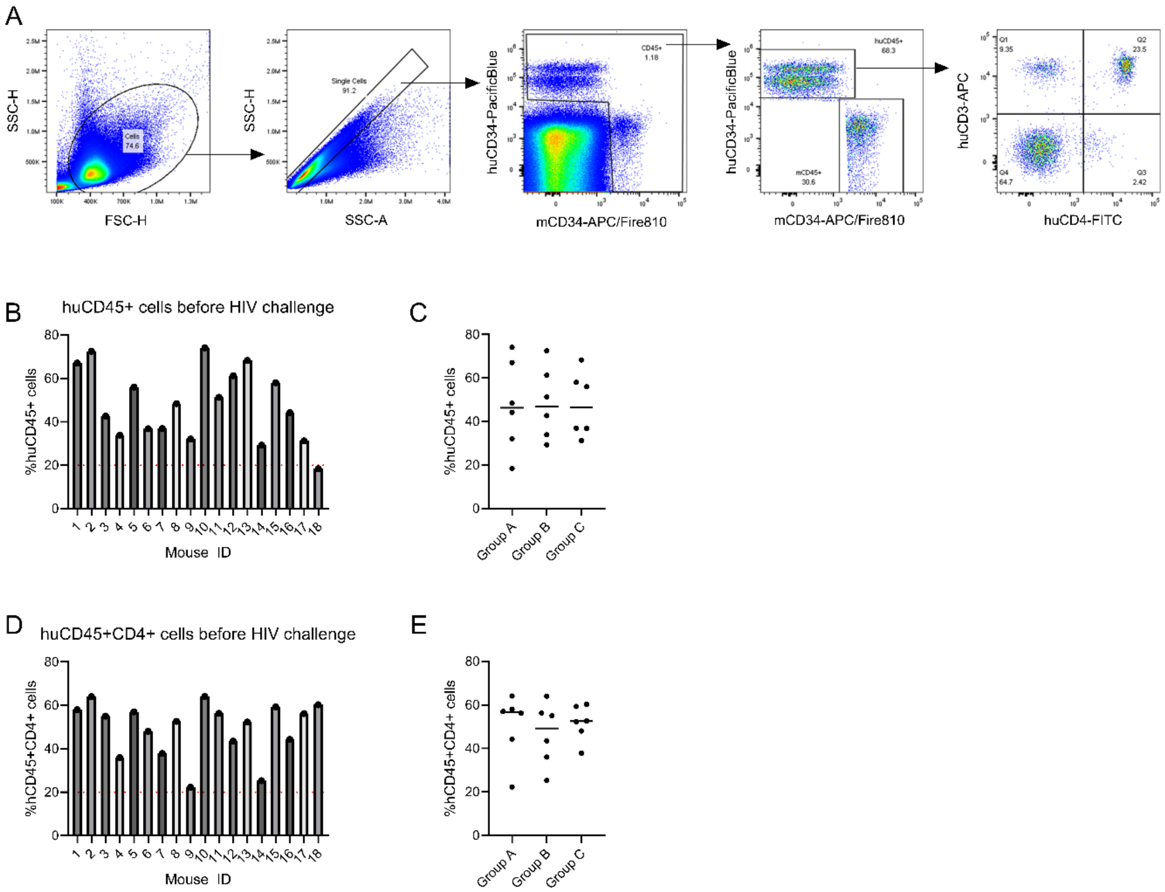


**Supplemental Figure 1. Method for assaying reconstitution of human immune cell population and sorting animals for experiments. A.** Gating strategy for Flow cytometry analysis to identify human immune cell populations in peripheral mouse blood. **B-E.** Example allocation of animals across treatment groups to achieve normalized huCD45$^+$ and huCD45$^+$huCD4$^+$ populations. Peripheral huCD45$^+$ population (**B**) and huCD45$^+$huCD4$^+$ populations (**D**) at 13 weeks post human stem cell transplant. Distribution of huCD45$^+$ population (**C**) and huCD45$^+$huCD4$^+$ populations (**E**) in mice across three treatment groups.

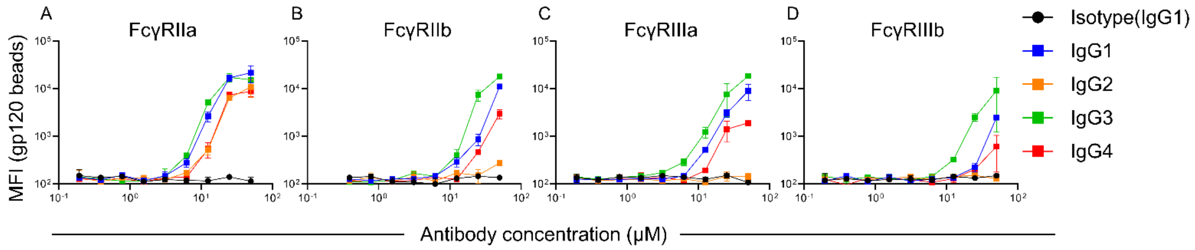


**Supplemental Figure 2. VRC01 IgG subclasses binding to human FcγR. A-D**. Multiplex bead-based binding profiles of VRC01 IgG1 subclasses to human receptor proteins FcγRIIa (**A**), FcγRIIb (**B**), FcγRIIIa (**C**), and FcγRIIIb (**D**).

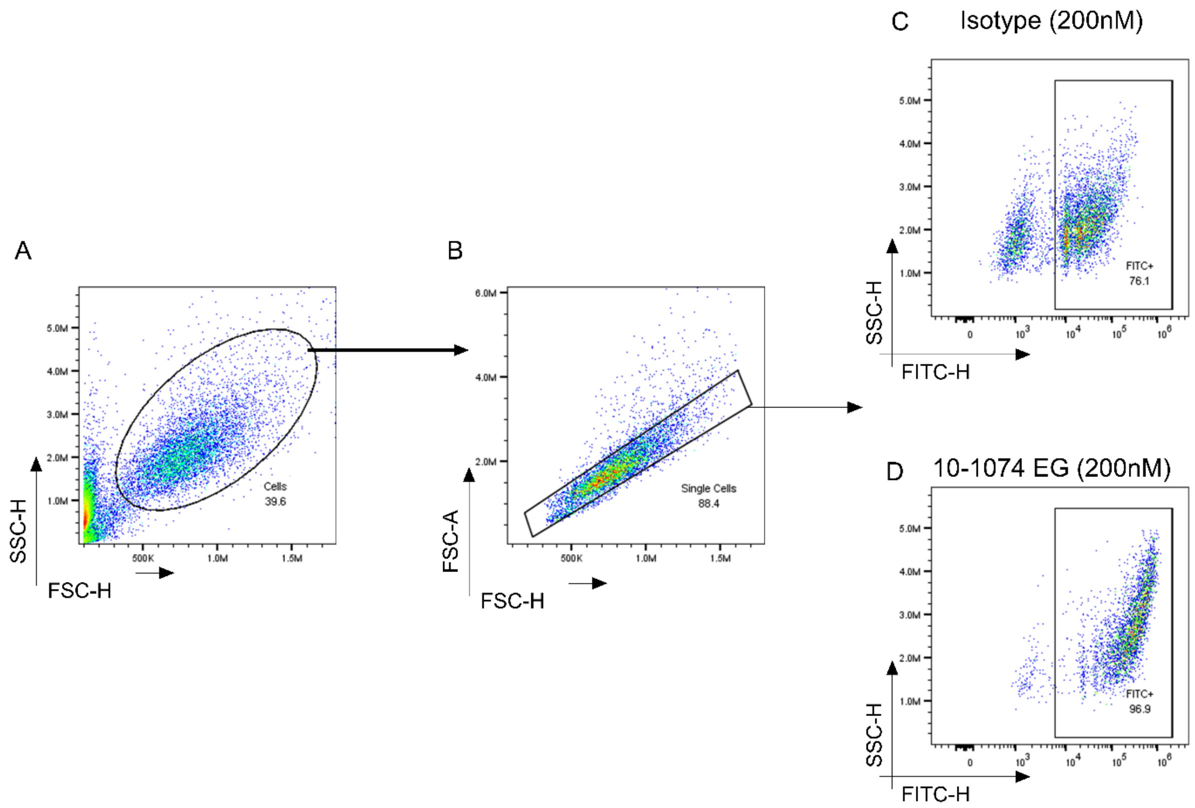


**Supplemental Figure 3. Gating strategy for assessing antibody-dependent cellular phagocytosis (ADCP) via THP-1 cell assay. A** representative example demonstrating gating strategy for A. selecting THP-1 cells, **B.** excluding doublets, to then **C-D.** measure FITC signal from phagocytosed gp120-bound fluorescent beads. Examples to demonstrate FITC signal in this assay when the sample has 200nM of either isotype control (**C**) or 10-1074 EG (**D**).
